# Simulation-based Reconstructed Diffusion unveils the effect of aging on protein diffusion in Escherichia coli

**DOI:** 10.1101/2023.04.10.536329

**Authors:** Luca Mantovanelli, Dmitrii S. Linnik, Michiel Punter, Hildeberto Jardón Kojakhmetov, Wojciech M. Śmigiel, Bert Poolman

## Abstract

We have developed Simulation-based Reconstructed Diffusion (SbRD) to determine diffusion coefficients corrected for confinement effects and for the bias introduced by two-dimensional models describing a three-dimensional motion. We validate the method on simulated diffusion data in three-dimensional cell-shaped compartments. We use SbRD, combined with a new cell detection method, to infer the diffusion coefficients of a set of native proteins in *Escherichia coli.* We observe slower diffusion at the cell poles than in the nucleoid region of exponentially growing cells. We find that this observation is independent of the presence of polysomes. Furthermore, we show that the newly formed pole of dividing cells exhibits a faster diffusion than the old one. We hypothesize that the observed slowdown at the cell poles is caused by the accumulation of aggregated or damaged proteins, and that the effect is asymmetric due to cell aging.

## Introduction

Diffusion of molecules inside cells plays a crucial role in the functioning of biochemical processes. In prokaryotic cells, which lack membrane-bound compartments, except for the periplasm in Gram­negative bacteria, the majority of cellular processes take place in the cytoplasm. Here, the random motion of (macro)molecules allows for the functioning of highly sophisticated systems, such as the correct localization of the septation ring in *Escherichia coli,* governed by a reaction-diffusion mechanism ^1^, the signal transduction that leads to chemotaxis ^2^, or the translation of mRNA into proteins by polysomes, ribosomes-mRNA assemblies, mostly localized at the cell poles ^3–6^.

The cytoplasm of bacteria is an extremely crowded environment, where the concentration of macromolecules, mainly proteins and RNAs, can reach values up to a volume fraction of 15-20% in growing cells ^7–9^ and even higher in osmotically stressed cells ^10–13^. The molecular composition of the cytoplasm is highly diverse, with molecules spanning in size over more than three orders of magnitudes, from sub-nanometric for ions and metabolites, to micrometric for the chromosome ^14, 15^. Despite this variation in size and surface properties of the molecules, many cellular components are uniformly distributed throughout the cell ^5, 16, 17^, with notable exceptions like the chromosome and nucleoid-binding proteins (localized in the cell center^4, 18^), the polysomes (localized at the poles and cytoplasmic periphery ^3, 4, 6^), and aggregated or misfolded proteins (localized at the cell poles^19–21^) (Fig. 1). In addition, there is increasing evidence for the formation of phase-separated liquid droplets or biomolecular condensates in the cytoplasm of microorganisms ^22–24^, which are metastable structures where certain proteins partition.

**Fig. 1.**
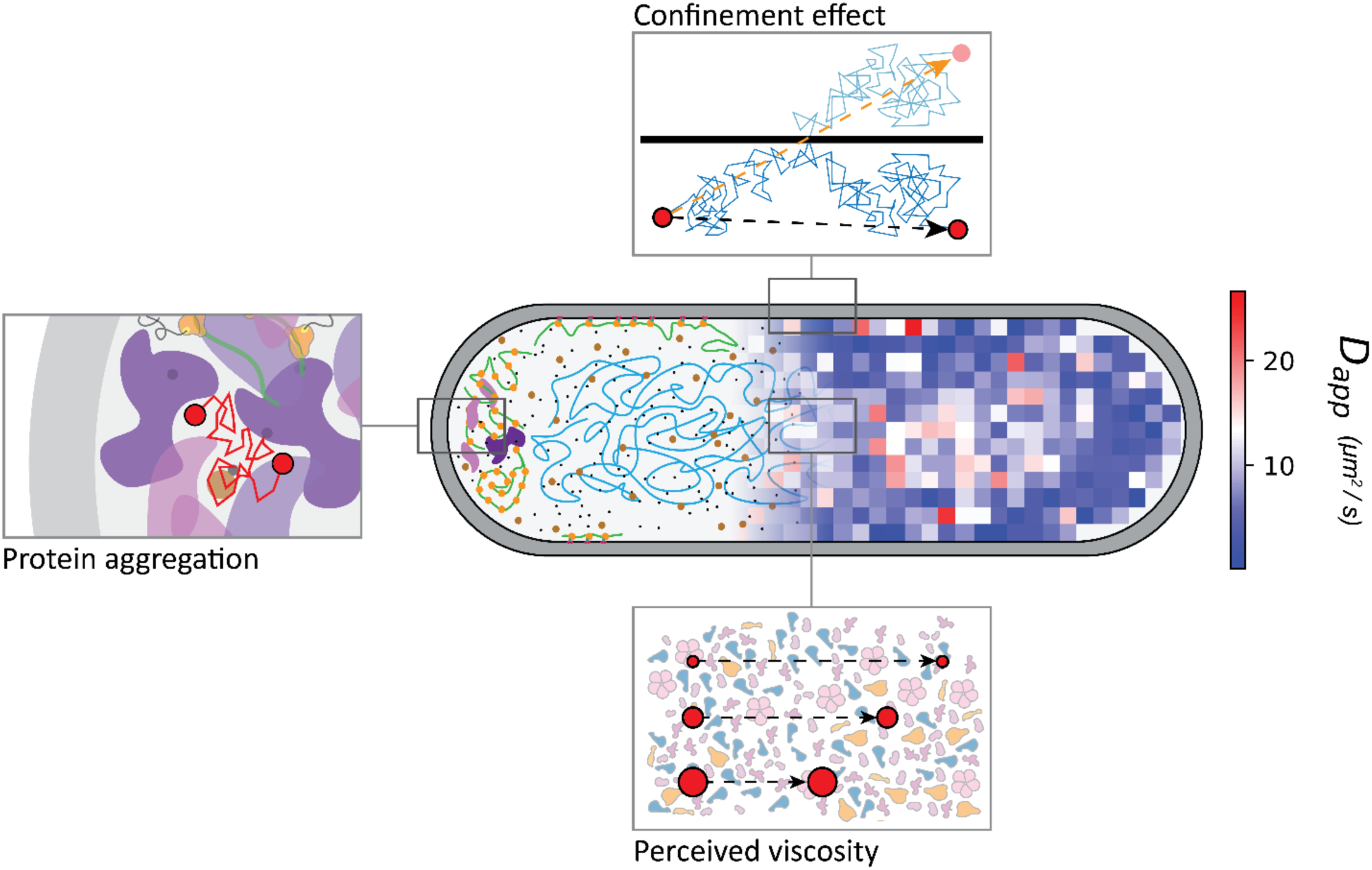
Diffusion in the cytoplasm of *E. coli.* A diffusion map obtained with SMdM is overlayed with a schematic of the cytoplasm of the cell. The top panel highlights the effect of confinement on the measured diffusion, which leads to lower diffusion coefficients near the boundaries of the cell. The bottom panel represents the effect of the perceived viscosity by diffusing proteins. Since diffusion scales with the complex mass, bigger particles will be affected more by the crowding of the cytoplasm than smaller molecules and move relatively more slowly, leading to the deviation from the Einstein-Stokes equation. The left panel represents our current hypothesis on the observed slowdown at the cell poles compared to the cell center, with accumulation of aggregated or misfolded proteins impairing the diffusion in these regions.

Diffusion of spherical particles in aqueous solutions can be described by the Einstein-Stokes equation ^25^. However, motion of particles in the highly crowded and inhomogeneous cytoplasm of bacterial cells has been extensively documented to deviate from the aforementioned model^16, 17, 26, 27^. We have shown that the apparent diffusion is solely dependent on the complex mass, that is the molecular weight of the monomer, summed with the molecular weight of the fluorescent reporter, multiplied by the oligomeric state of the analyzed molecules ^16^ (Fig. 1). The same conclusion was recently obtained in another study, using a different method ^17^. Apparent diffusion of proteins interacting with large cellular components, such as the cell membrane, the nucleoid, the ribosome or the proteasome complex, depends on the sum of both the interacting masses and on the interaction strength (dissociation constant, K_D_) of the species ^17, 28^. The deviation from the Einstein-Stokes equation indicates that diffusion in cells depends not only on the size of the analyzed molecules, but also on the composition and physical state of the cytoplasm. This, in turn, is dependent of e.g. (fluidization by) metabolism ^29, 30^, catalysis-induced enzyme movement^31, 32^ and environmental stresses ^11, 33^. The deviation from Stokes-Einstein can be explained by taking into account both the solvent quality of the cytoplasm and the size of the diffusing species. The macromolecular crowding experienced by each protein is dependent of its own molecular weight, that is, smaller molecules will be less affected by the crowded cytoplasmic environment than bigger ones ^16, 17, 26, 27, 34^ (Fig. 1). We have shown that this so-called perceived macromolecular viscosity is not spatially uniform in the cell^16^.

Despite the advancements made in the field of single molecule fluorescence microscopy ^16, 17, 35–39^, diffusion measurements are still highly influenced by the effect of confinement, especially in small compartments such as the bacterial cytoplasm ^16^ and periplasm ^40, 41^, and eukaryotic organelles ^42^. Here, diffusion coefficients near the cell boundaries always appear lower than in the cell center ^16, 38^. This does not allow to properly separate the effect of confinement from possible physiological slowdown in diffusion of the analyzed species ^16^ (Fig. 1). Moreover, techniques such as Single Particle Tracking (SPT) and Single Molecule displacement Mapping (SMdM) produce a two-dimensional output of a three- dimensional motion, which leads to obvious shortcomings in the estimation of diffusion coefficients of particles freely moving in the cell cytoplasm.

Some methods to resolve confined diffusion have been developed. Bickel ^43^ proposed a mathematical method to obtain the mean square displacement in disks and sphere for particles. Bellotto et al. ^17^ derived a Ornstein-Uhlenbeck model for fitting Fluorescence Correlation Spectroscopy (FCS) data acquired in a confined cylinder of infinite length, but not considering the effect of the cell poles, which represent the zones where the confinement affects the observed diffusion the most^16^.

A detailed analysis of the diffusion of macromolecules in the cell poles is paramount to understand the effects of aging in rod shaped cells. A study that followed repeated divisions of *E. coli* suggests that cells that inherit the old pole exhibit a diminished growth rate, decreased offspring production, and an increased incidence of death ^44^. Asymmetry in the doubling time of old and new pole daughter cells has also been observed in a more recent study in *E. coli*^45^. The underlying mechanisms of aging of bacteria are at best poorly understood.

In this study, we set out to solve the shortcomings of previous methods for analyzing confined diffusion in small compartments, and we apply the new tools to investigate the relationship between diffusion in the cell pole regions and aging. We developed a method to determine diffusion coefficients that are not affected by the effect of confinement or motion along the third dimension. We developed the technique to unveil novel properties of diffusion in small compartments and we demonstrate its validity in *E. coli* cells. We first investigate a mathematical approach, followed by a simulation-based analysis, which we named Simulation-based Reconstructed Diffusion (SbRD). We highlight the limitation of using mathematical models to solve the problem of confinement, and the advantages of a simulation-based approach. We use the method, in combination with a new cell detection tool, to re-analyze a previously acquired dataset^16^ and obtain confinement-corrected values for the diffusion of molecules in the *E. coli* cytoplasm. Further, we investigate diffusion near the cell boundaries and at the cell poles to determine how much confinement influences the slowdown in these regions. We test the effect of antibiotics that disrupt the polysome structure in the apparent slowdown of diffusion at the cell poles. Finally, we make observations about asymmetry in diffusion in the bacterial cell, which we associate with aging.

## Results

### A mathematical solution to the limitations of confined diffusion

We previously showed through simulations that the measured diffusion coefficient is underestimated when particles move by random motion in a confined environment. The deviation from the real diffusion coefficient drastically increases when the diffusion coefficient gets higher, and the analyzed molecules are in a region closer to the boundary of the confinement^16^ (Fig. 2A).

**Fig. 2.**
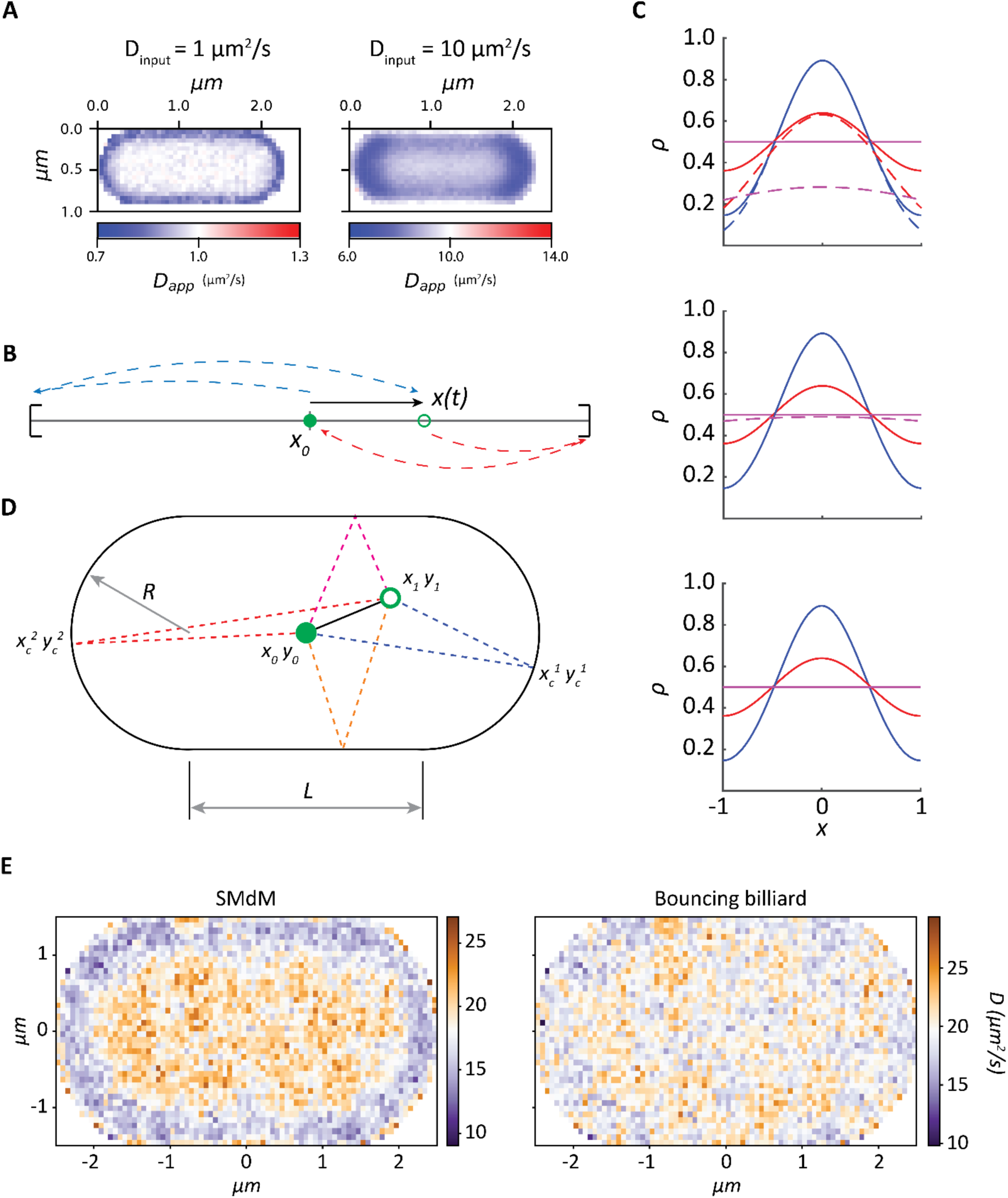
Mathematical solution to confinement. **(A)** Effect of confinement on diffusion. Diffusion simulations performed in a billiard at different input diffusion coefficients. The position of the particles was measured every 1.5 ms. The higher the diffusion coefficient used for the simulation, the more pronounced the confinement effect is. (B). Sketch of the (random) motion of a particle in a 1 dimensional closed interval. The point x_0_ stands for the initial position of the particle at t = 0. The point x(t) represents the position of the particle at time t > 0. In this scenario, and for fixed t > 0, there are several possibilities for measuring a particle’s position at x(t). The first scenario is that the particle travels to its measured position without bouncing. Another scenario is that the particle arrives at its measured position after bouncing (on the boundaries) once. In this way, there are infinitely many ways in which the particle can reach its measured position, depending on the number of bounces the particle made. However, the probability of each case is inversely proportional to the total distance traveled. In B we show the distances traveled for 0 and 1 bounce (against each boundary). (C) Solutions of the diffusion equation on a bounded interval with length *L = 2* µm, diffusion coefficient D = 2 µm^2^/s, and time *t =* {0.05, 0.1, 0.5} seconds, shown in blue, red, and purple respectively. The solid lines correspond to the analytical solution (Eq. 3), while the dashed curves correspond to (Eq. 5). From top to bottom we show comparisons for 0, 1, and 2 bounces. Notice that for this example, accounting for two bounces already gives a sufficiently good approximation of the analytical solution (bottom panel). (D) A few trajectories of a billiard in the Bunimovich stadium. We show one 0-bounce (solid line) and four 1-bounce billiard trajectories (dashed lines). The bouncing points 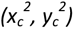 and 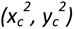 are solutions of the system of equations shown under (7). (E) Diffusion maps of a billiard obtained by analyzing a Smoldyn simulation created with an input diffusion coefficient of 20 µm^2^/s, with particles’ position measured every 1.5 ms. Maps are obtained via SMdM analysis (left) and via mathematical method analysis (right).

We now developed a mathematical model to solve this limitation. The simplest approach to modeling of diffusion is through Ficks laws, which in his second law led to the so-called diffusion equation (Eq. 1)

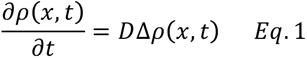

In the context of random motion, say a random walk of a particle, *p* denotes a probability density (or distribution) and 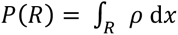 would be the probability of finding the particle within a region *R*.

Here we solve analytically the one-dimensional diffusion equation with reflecting boundaries. For simplicity, we interpret *ρ(x, t)* as the probability density for finding a particle at position x at time *t.* Assuming that at *t* = *0* the particle is located at x = 0, and that the probability density *ρ(x, t)* = 0 as x approaches infinite for any finite time, a solution to equation 1 is then given by equation 2:

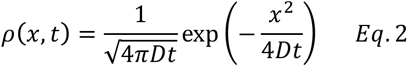

In a heuristic way, equation 2 provides a measure of the likelihood of finding a particle at position x after time *t* provided that the medium where it moves is infinite.

Let us now consider the diffusion of a particle within a bounded domain, for simplicity from - *L/2* to + *L/2,* where L is the length of the domain. We assume that the particle is located at x = 0 at *t* = 0, and that it is reflected back to the interior of the interval once it reaches the boundary (Fig. 2B). The analytical solution for this case is (see Supplementary Information - Diffusion on a closed interval):

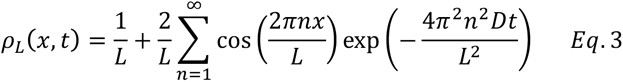

Relying on an analytical solution of the diffusion equation in higher dimensions and for complicated geometries is not convenient. Equation 2 is valid for diffusion in unbounded domains. We can approximate the solution of equation 3 using a “folding approach“, by accounting for the bounces a particle makes when hitting the boundaries, assuming no loss of energy in the process, and adding them up (Fig. 2C).

In this way, an approximation of equation 3 is given by equation 4:

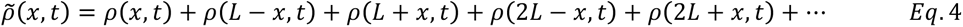

where the term *ρ(kL -x) + ρ(kL + x)* corresponds to the density for the particle being at x at time *t* after *k* bounces. When substituting equation 2 in equation 4 we get:

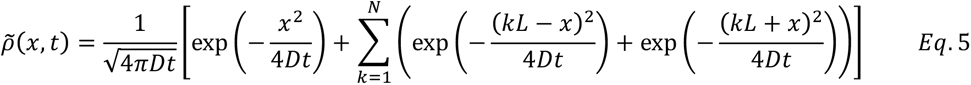

where *N* denotes the maximum number of bounces (Fig. 2C).

One can take a similar approach in higher dimensions. For example, let us consider the diffusion equation in a rectangular plate of sides *A* (horizontal) and *B* (vertical), and let us assume that the motion of the particle on each coordinate *(x, y)* is independent of each other. Then, the analytical solution for the diffusion equation is:

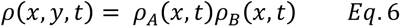

where *ρ_A_* and *ρ_B_* are as in equation 3.

We can however take another approach, as we did for the interval in one dimension (equation 4, 5). We denote by *p(t) = (x(t), y(t))* the position of a particle in the rectangular plate at time *t.* As above, we can compute the density *ρ(x, y, t)* by adding up all the densities corresponding to trajectories in the rectangular plate that take the particle from the initial position *p_0_* to some final position *p_f_* after a number of bounces (Supplementary fig. 1A, 1B). What we describe is an example of a mathematical billiard.

To use these ideas to estimate the diffusion coefficient inside a cell, we approximate the geometry of the cell by a planar sphero-cylinder, also known as a Bunimovich stadium (Fig. 2D). In this setting, 0-bounce and 1-bounce trajectories can be easily computed. Densities corresponding to trajectories that bounce on the straight (top and bottom) sides of the sphero-cylinder are computed as aforementioned. Those corresponding to bounces on the circular sides can be computed by solving the system of equations:

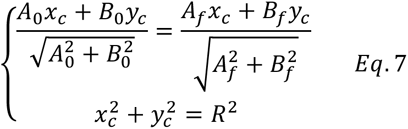

where *A_0_ = y_c_ – y_0_, B_0_ = x_c_ – x_0_, A_f_ = y_c_ – y_f_, B_f_ = x_c_ – x_f_,* and *(x_0_, y_0_), (x_c_, y_c_), (x_f_, y_f_)* represent the starting point, a bouncing point on the circular section of the cell boundary and the end point, respectively (Fig. 2D).

Since equation 7 cannot be solved analytically, we developed a script (see Supplementary Material - algorithm 1) to compute the 0-bounce and 1-bounce trajectories of any diffusing particle, provided that the start and end positions are known. We then used this algorithm to calculate the diffusion coefficient of two-dimensional diffusion simulations in a billiard, generated with Smoldyn ^46^ (Fig. 2E). Smoldyn allows simulation of the motion of particles using a predefined diffusion coefficient and time resolution, within a simulation compartment. With our mathematical model we obtain a final diffusion coefficient for every pixel that is a good approximation of the input diffusion coefficient used for the simulations.

## Limitations of a mathematical approach

Applying the mathematical approach to solve the diffusion equation near the boundaries in confined environments has three shortcomings. (i) The approximation (equation 5) is valid for the given boundary conditions, but not for e.g. non-continuous, non-convex surfaces. An example of a surface where our model would have failed is the Penrose unilluminable room ^47^ (Fig. 3A), or the matrix of mitochondria. Some regions of these surfaces are inaccessible by rays that start from particular locations, regardless of the number of bounces. However, for a particle freely diffusing in any compartment, it would be possible to reach any location, leading to the emergence of starting and final points that cannot be connected by reflecting rays. (ii) The model cannot be extended from the two-dimensional billiard to the three-dimensional spherocylinder. Given a start and end point in two dimensions, it is always possible to find a reflection point on a circle; on the other hand, given a start and an end point in three dimensions, there will be an infinite number of reflection points on a sphere. Therefore our model implies that the motion of particles only occurs in two dimensions *(x,y* coordinates of the diffusing particles), while in reality (in cells) particles also diffuse along the *z*-axis. When simulating diffusion in a three-dimensional spherocylinder and analyzing it with our mathematical model (equation 5), we observed an underestimation of the diffusion coefficient throughout most of the cell, and an overestimation of it close to the boundary (Fig. 3C). The underestimation is due to the motion along the *z*-axis, which is not accounted for in our model, leading to a measured step length shorter than the actual one (Fig. 3B, left). The overestimation is likely due to particles bouncing against the boundary at *z* coordinates where the spherocylinder *(x,y)* section has a smaller perimeter (Fig. 3B, right). (iii) Finally, when approximating the solution in a bounded domain one must compute all trajectories that lead from the initial position p_0_ to the final position p_f_ after *0, 1, 2, …, N* bounces. Computing all these trajectories analytically is cumbersome, and therefore we limited our analysis to the 1-bounce case. This can be limiting in the case of fast diffusion in small compartments: when the square root of the mean square displacement is much higher than the size of the compartment, or when the acquisition time is very long, a particle could bounce against the surface multiple times over the acquisition period.

**Fig. 3.**
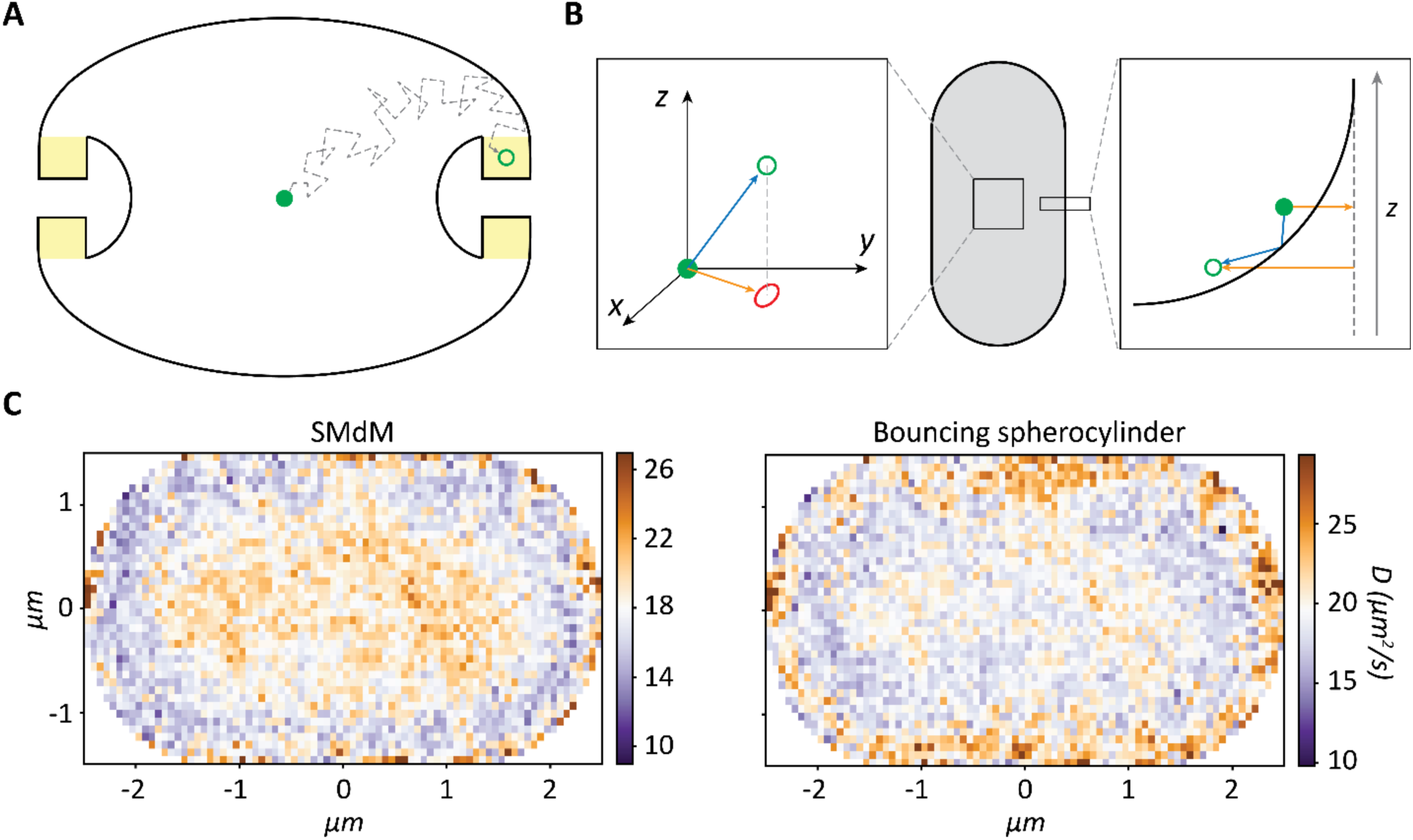
Limitation of the mathematical model for confined diffusion. **(A)** Schematic of Penrose unilluminable room. A ray (vector) starting from the center of the room can never reach the regions colored in yellow, regardless of the number of bounces against the perimeter. A particle moving by random motion (dashed line) can reach any region in the room. (B) Limitations of using a 2D model to describe a 3D motion. Left panel: the motion of a particle moving in 3D space (blue arrow) is projected on a 2D surface (orange arrow). The observed distance is shorter than the actual travelled distance, leading to an underestimation of the diffusion coefficient. Right panel: the effect of the overestimation of the billiard perimeter. By observing the projection of a 3D spherocylinder in 2D, we use as billiard’s perimeter its largest *×y* projection. The bouncing of particles will likely occur against a different section of the spherocylinder, where the circumference of the billiard is smaller, when displacements are binned in a pixel close to the boundaries, assuming the starting points of such displacements are in *z-stacks* other than the section of the spherocylinder with the largest circumference (*z = 0* for a spherocylinder centered *in (0,0,0)*). In this way, the calculated bouncing path (orange arrow) overestimates the actual path (blue arrow), leading to an overestimation of the diffusion coefficient near the boundaries. (C) Diffusion maps of a spherocylinder obtained by analyzing a Smoldyn simulation created with an input diffusion coefficient of 20 µm^2^/s. Maps are obtained via SMdM analysis (left) and via mathematical method analysis (right).

Despite these caveats, we have shown and verified that the proposed “folding approach“ is reasonable and, in fact, the shortcoming identified above are merely computational: (i) Complicated geometries can be approximated by the union of convex sets, similar to a triangulation of smooth objects. For each convex set one can potentially tune and adapt our “folding“ approach, (ii) In three dimensions one can extend the description for the rectangular plane to diffusion in cubes. Consequently, if the three dimensional confined geometry can be approximated by triangulated cubes, one can still apply our “folding“ argument, (iii) Analytic computation of trajectories is computationally expensive. However, we have shown that based on a few bounces, the estimation of the diffusion coefficient already improves significantly. If one pairs this with an approximation of complicated geometries by simpler ones, one can still make sufficiently good approximations.

### A simulation-based solution to the limitations of confined diffusion

We then reasoned that an approach based on diffusion simulations could have advantages compared to mathematical models, as the shape of the compartment or the number of bounces against the surface would not be limiting for the outcome. In brief, we developed a method to recursively estimate the diffusion coefficient with an algorithm that makes use of Smoldyn ^46^ and the SMdM technique ^16, 35^ (Fig. 4A). Smoldyn allows the simulation of the motion of particles within a compartment. The compartment can be either mathematically described, or an input of triangulated coordinates of any desired shape. Confinement is accounted for in the motion of the particles, which reflect off of the boundaries without losing velocity. We generated diffusion simulations with Smoldyn inside a spherocylinder, and as anticipated the apparent diffusion is underestimated, especially in regions close to the boundaries (Fig. 2A, 2E, 3C). The extent of the underestimation is proportional to the diffusion coefficient, and inversely proportional to the size of the confined space and to the acquisition time. Our algorithm yields a diffusion coefficient that is homogenous throughout the whole compartment and matches the predefined value used to create the diffusion simulations.

**Fig. 4.**
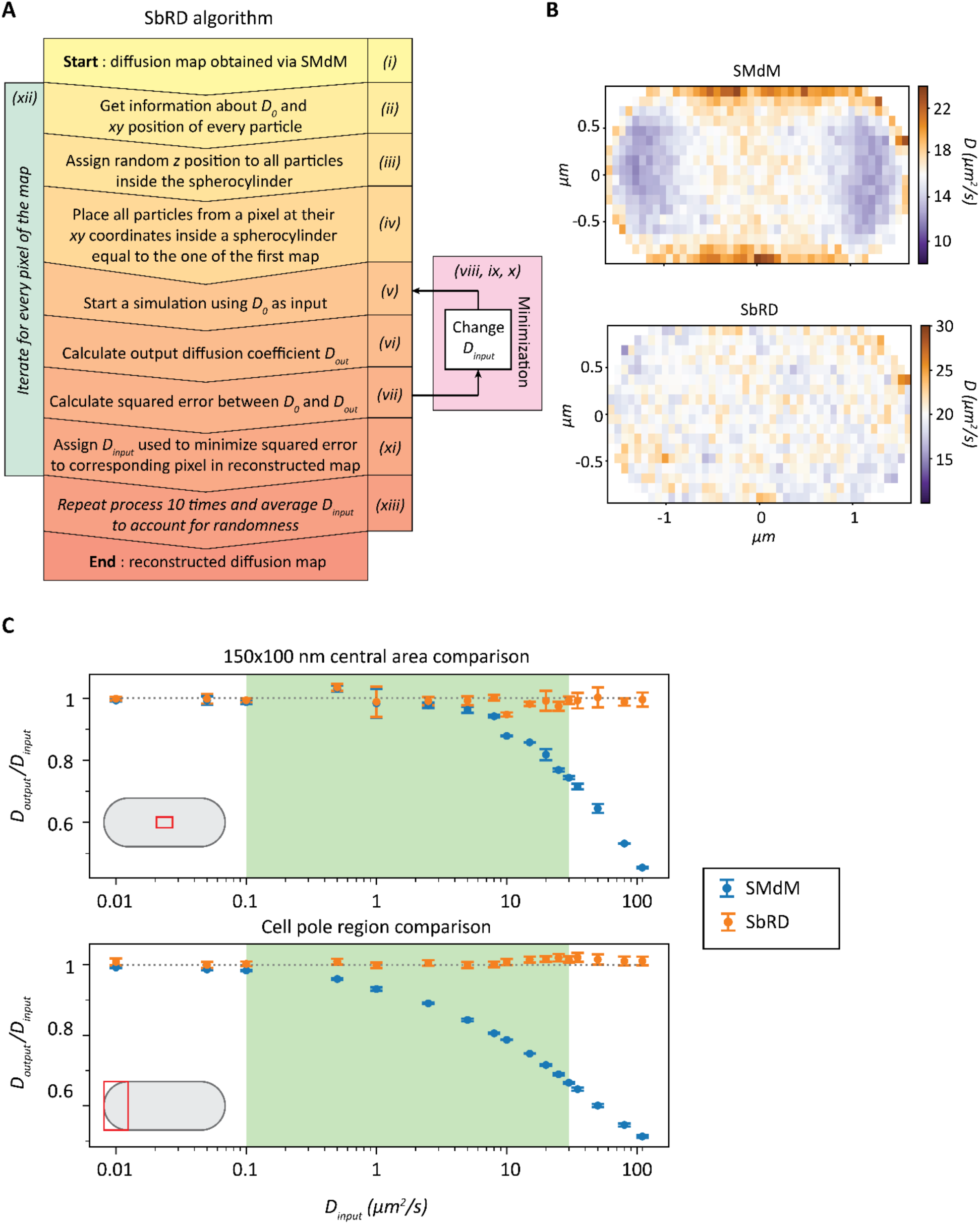
Simulation-based Reconstructed Diffusion. **(A)** Algorithm representation of SbRD. **(B)** Diffusion maps of a spherocylinder obtained by analyzing a Smoldyn simulation created with an input diffusion coefficient of 20 µm^2^/s. Maps are obtained via SMdM analysis (top) and via SbRD (bottom). **(C)** Comparison of the dependence of the ratio of *D_output_ / D_input_* in a simulated spherocylinder when analyzing the centermost 100 nm by 150 nm area (top) and the cell pole area (bottom) with SMdM and SbRD. The gray dotted line represents the ideal case in which the ratio of *D_output_* over *D_input_* is one. The relevant range of diffusion coefficients for proteins in the cytoplasm is highlighted in green.

The main steps for the operation of the algorithm are: (i) For simulations, an input diffusion coefficient (*D_input_*^0^) is used to simulate the diffusion of particles in a spherocylinder, as previously described ^16^. For *in vivo* datasets, diffusing particles are measured via stroboscopic illumination microscopy, as previously described. The diffusion map is experimentally obtained by SMdM ^16, 35^; (ii) The diffusion map yields the total number of displacements per cell and the measured diffusion coefficient (*D_output_*^0^) for every position; (iii) The starting (x,y) coordinates of all the observed particles are used to place them inside a simulated spherocylinder, and their *z* coordinates are randomly assigned. This is done by taking a value from a uniform random distribution ranging from the lowest to the highest *z* value that a particle can have at that specific (x,y) position inside the spherocylinder; (iv) The starting positions (*x,y,z*) of the particles that belong to a specific pixel of the original diffusion map are selected; (v) Diffusion simulations are run, using as input diffusion coefficient the *D_output_°* value obtained via SMdM; (vi) The diffusion observed by SMdM is analyzed to obtain a new diffusion coefficient for the specific pixel (*D_output_*^1^); (vii) The absolute difference (*ε*) between *D_output_*^0^ and *D_output_*^1^ is calculated; (viii) The diffusion simulation process (and subsequent SMdM analysis) is repeated recursively, every time using a different input diffusion coefficient (*D_input_^i^*). (ix) The difference (ε) between *D_output_°* and the one obtained through the last simulation (*D_output_^i+1^*) is calculated; (x) The recursion is run until the difference between *D_output_^N^* and *D_output_°* reaches a minimum; (xi) The input value (*D_input_^N-1^*) that led to the minimal difference between *D_output_^N^* and *D_output_*^0^ is then used to create a new map, which carries *D_input_^N-1^* in the position of the analyzed pixel; (xii) The process is repeated for every pixel, until a map is obtained in which the diffusion coefficients represents the unbiased diffusion values, that is diffusion values that are corrected for the effect of confinement and for the underestimation caused by a two-dimensional representation of a three-dimensional motion. (xiii) The whole routine is repeated ten times for each cell. The output maps obtained in each iteration are averaged to account for the randomness introduced when assigning the *z* starting position to all particles and for the randomness introduced by Smoldyn in simulating the diffusion of particles ^46^. (Fig. 4A).

### Simulation-based Reconstructed Diffusion overcomes limitations of confinement caused by cell boundaries

Simulation-based Reconstructed Diffusion (SbRD) together with SMdM allows obtaining more accurate diffusion maps. It is possible to retrieve the actual diffusion coefficient also for the regions close to the cell boundaries (Fig. 4B). We also simulated scenarios of a cell displaying slower diffusion at one cell pole, and observed that SbRD is correctly identifying the region with slower diffusion and higher diffusion coefficients in the rest of the cell (Supplementary Fig. 2).

We then benchmarked SbRD against SMdM by varying *D_input_* from 0.01 to 110 µm^2^/s and analyzing *D_output_* in the innermost 100 nm^2^ square region of the simulated spherocylinder, which is the region least affected by the effect of confinement, and in a cell pole, which is the region most affected by the effect of confinement. We observe for the innermost region that *D_output_* obtained via SMdM decreases to 90% of its input value already for *D_input_* of 10 µm^2^/s, and that the deviation increases as expected with *D_input_* (Fig. 4C, top panel). The decrease is more pronounced when the cell pole is analyzed (Fig. 4C, bottom panel), with *D_output_* obtained via SMdM decreasing to 90% of its input value already for *D_input_* of 1 µm^2^/s. Importantly, the output of SbRD remains stable throughout the whole set of measurements, with diffusion coefficients near 100% of *D_input_* (Fig. 4C).

### Billiard fitting of rod-shaped bacteria

The shape of *E. coli* cells is generally assumed to be a spherocylinder ^48–52^. Software is available to determine the shape of *E. coli* cells from two-dimensional images ^53–55^, but estimation and subsequent triangularization of their three-dimensional shape does not necessarily lead to the true shape of the cell. For instance, invaginations or protuberances observed on the *xy* plane might not be observed on the *xz* plane. By analyzing our microscopy data, we observed that the vast majority of cells had a shape that could be very well approximated by a two-dimensional projection of a spherocylinder. Therefore, we decided to use this shape for cell detection and modeling, which at the same time allowed for an easy implementation in Smoldyn. To apply SbRD to microscopy data, we developed an algorithm to automatically detect cells as billiards in microscopy images (Fig. 5A). Firstly, each held of view was filtered for background noise, yielding clusters of points representing cells. Point clouds were rotated so that their major axis was aligned to the x-axis, and subsequently clustered using the equation of a billiard (Eq. 8).

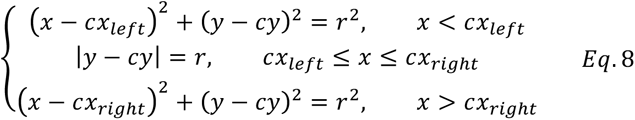

**Fig. 5.**
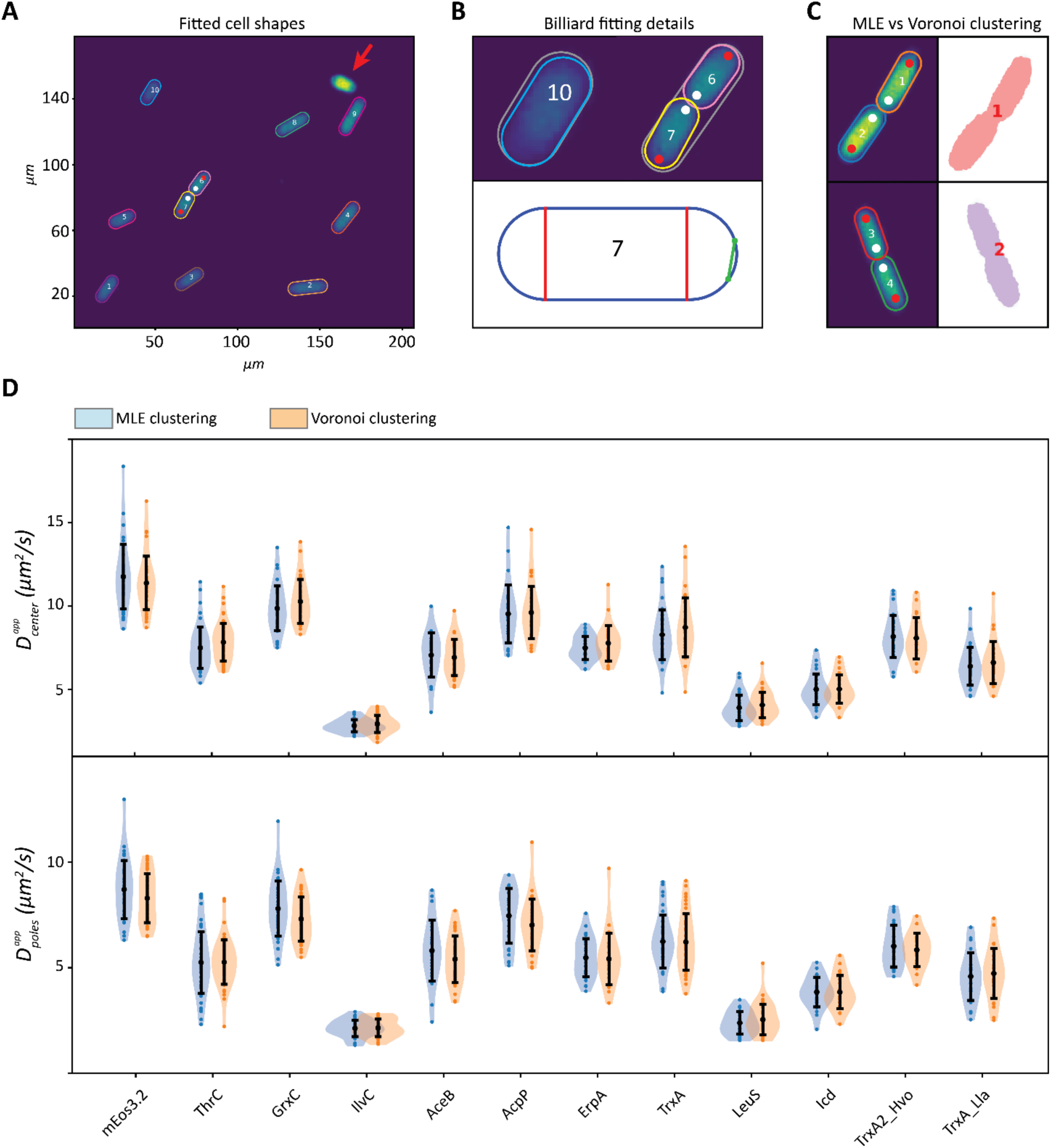
Maximum likelihood-based detection method. **(A)** Field of view and fitting of billiards around the identified cells, using a maximum likelihood estimation method. The cell indicated by a red arrow was discarded because it has too many displacements. (B) Details of the fitting process. In the top panel, the initial guess used for fitting, encompassing all the points clustered as a single cell, is represented in grey, while the final fitting is colored. For cells that just completed division, the initial guess encompasses both cells. Due to the abnormal length of the cell, the fitting routine is automatically performed with two billiards that detect both cells. Using the fitting information it is possible to identify the newly formed cell pole (white dots) and the old one (red dots). The bottom panel shows the billiard used to describe the shape of cell 7. Since cells are represented as billiards, it is possible to obtain accurate estimates of their length and radius, which allow distinguishing the cell poles and cell center for every cell. For cells that just divided and two billiards overlapping, the intersection points are calculated (green dots) and used to draw a line (green line), which is then used to properly model the spherocylinder in the SbRD routine. **(C)** Comparison of Maximum likelihood method and Voronoi clustering for cell detection. Voronoi clustering cannot properly distinguish cells that are too close to each other. (D) Comparison of the apparent diffusion coefficients obtained with SMdM by analyzing the central region (top) and the poles (bottom) of cells identified with Voronoi clustering and with our maximum likelihood method, from images acquired in our previous work^16^. Curves are obtained via kernel density estimation.

We refine the cell selections via Maximum Likelihood Estimation using the following assumptions (see Methods - cell clustering and detection): (i) fluorescent points are uniformly distributed throughout the cell; (ii) the cells, here *E. coli,* are modeled as spherocylinders; therefore the 2D projection of their fluorescence (shape of a billiard) appears more populated in the center than near the cell boundary; (iii) all the fluorescent spots observed in a cell have the same probability of being noise; and (iv) every fluorescent point is equally likely to be noise or to be a photoconverted mEos3.2. We then filtered the selected cells for a minimal length of 0.65 µm and a maximal width of 1.5 µm (see methods - cell clustering and detection). If the length of the final billiard describing the shape of the cell was bigger than 3 µm, the algorithm separates the cluster into two billiards, each having half the length of the original one. The refinement step was then repeated for every cluster of two-billiards. This allowed the accurate detection of newly divided cells (Fig. 5B), which is often not possible with standard clustering methods, such as Voronoi (Fig. 5C). In our previous work ^16^ we acquired a dataset of diffusing proteins of different complex mass (Supplementary Table 1) using SMdM. Here, each cell was clustered using Voronoi clustering. We re-analyzed the full dataset from our previous work with the new clustering method. Importantly, we observe no significant difference between the two datasets, both in the number of detected cells and in the diffusion coefficients obtained via SMdM analyses for the cell center (Fig. 5D, top) and for the cell poles (Fig. 5D, bottom). We then used the information of each cluster to recreate a spherocylinder in Smoldyn ^46^ having the shape of the corresponding cell, to which we applied the SbRD algorithm (see “A simulation-based solution to the limitation of confined diffusion“ from point iii to point xi) to reconstruct diffusion maps, corrected for the confinement effect and bias by 2D models to describe a 3D motion, of the single-molecule fluorescent microscopy data.

### SbRD correlates the confinement-corrected diffusion coefficient with the perceived viscosity of the cytoplasm

An advantage of our clustering method is the possibility to precisely identify the cell poles and the cell center by using the radius of the billiard (Fig. 5B), allowing us to analyze these regions separately for every cell. We then compared the results obtained via SbRD with the results obtained via SMdM on our previous dataset ^16^ (Supplementary Table 1), and using the new clustering method for cell detection (Table 1, Fig. 6). We observe a significant difference in the observed diffusion values for faster diffusing proteins, while for the slower diffusing particles the difference appears to be not significant. Notably, we observe a higher statistical significance in the difference between the diffusion coefficient observed at cell poles. These results are in line with the observations made via simulations, which indicate that (i) given the fast acquisition time used in SMdM, the effect of confinement on the measured diffusion results less pronounced for slower diffusing particles; and (ii) that the confinement effect is more pronounced in the cell poles than in the cell center.

**Fig. 6.**
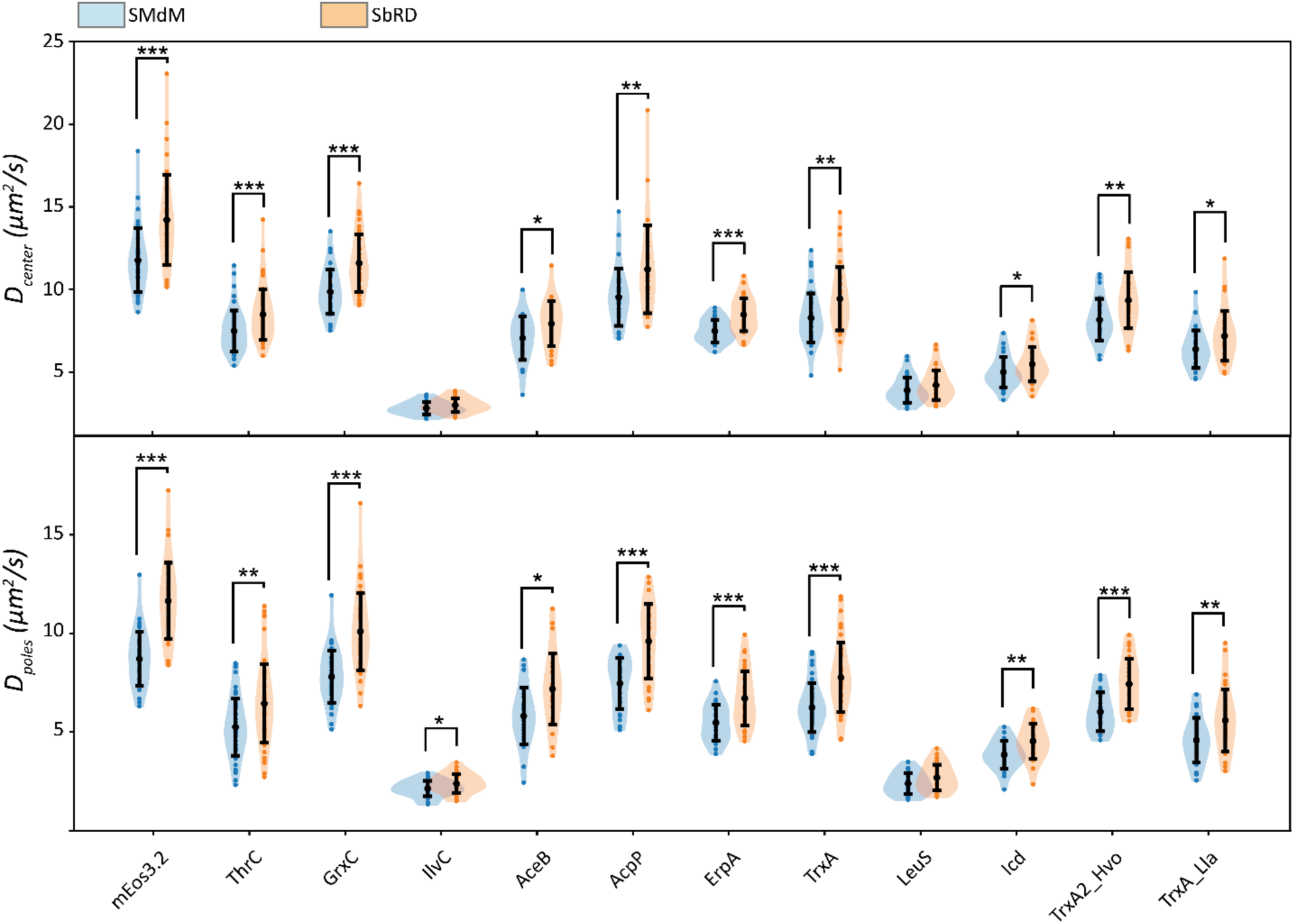
Comparison of the diffusion values obtained via SMdm and via SbRD. Comparison of the apparent diffusion coefficient obtained via SMdM and of the confinement-corrected diffusion coefficient for both the cell center (top) and the cell poles (bottom) for the dataset of proteins tagged with mEos3.2 ^16^. Asterisks indicate statistical significance obtained via a Mann-Whitney U test for non-normally distributed data.

**Table 1.** Lateral diffusion coefficients of cell center and cell poles obtained via SMdM and SbRD for constructs fused to mEos3.2. The columns show the name of the protein, their complex mass, their diffusion values obtained via SMdM (Dapp) for cell center and cell poles, and the confinement-corrected diffusion values obtained via SbRD (Dec). The Uniprot ID is provided for every protein, except for mEos3.2, for which the Fpbase ID is given.

| Protein name | UNIPROT ID | complex mass (kDa) | $D_{\text{app}}^{\text{center}} (\mu\text{m}^2/\text{s})$ | | $D_{\text{app}}^{\text{poles}} (\mu\text{m}^2/\text{s})$ | | $D_{\text{cc}}^{\text{center}} (\mu\text{m}^2/\text{s})$ | | $D_{\text{cc}}^{\text{poles}} (\mu\text{m}^2/\text{s})$ | |
| --- | --- | --- | --- | --- | --- | --- | --- | --- | --- | --- |
|  |  |  | mean | standard deviation | mean | standard deviation | mean | standard deviation | mean | standard deviation |
| mEos3.2 | VUXFR* | 25.7 | 11.8 | 1.9 | 8.7 | 1.4 | 14.2 | 2.7 | 11.6 | 1.9 |
| ThrC | P00934 | 72.8 | 7.5 | 1.2 | 5.3 | 1.5 | 8.5 | 1.5 | 6.4 | 2.0 |
| GrxC | 90AC62 | 34.8 | 9.9 | 1.3 | 7.8 | 1.3 | 11.6 | 1.7 | 10.1 | 2.0 |
| IlvC | P05793 | 318.9 | 2.8 | 0.4 | 2.1 | 0.4 | 3.0 | 0.4 | 2.4 | 0.5 |
| AceB | P08997 | 85.9 | 7.0 | 1.3 | 5.8 | 1.4 | 7.9 | 1.4 | 7.2 | 1.8 |
| AcpP | P0A6A8 | 34.3 | 9.5 | 1.7 | 7.5 | 1.3 | 11.2 | 2.7 | 9.6 | 1.9 |
| ErpA | P0ACC3 | 75.5 | 7.5 | 0.7 | 5.5 | 0.9 | 8.5 | 1.0 | 6.7 | 1.4 |
| TrxA | P0AA25 | 37.5 | 8.3 | 1.5 | 6.2 | 1.3 | 9.4 | 1.9 | 7.8 | 1.8 |
| LeuS | P07813 | 122.9 | 3.9 | 0.8 | 2.4 | 0.5 | 4.2 | 0.9 | 2.7 | 0.7 |
| Icd | P08200 | 142.8 | 5.0 | 0.9 | 3.8 | 0.7 | 5.5 | 1.1 | 4.5 | 0.9 |
| TrxA2_hvo | A0A558GCJ2 | 37.8 | 8.2 | 1.3 | 6.0 | 1.0 | 9.3 | 1.7 | 7.4 | 1.3 |
| TrxA_Lla | A0A089XQE8 | 37.4 | 6.4 | 1.1 | 4.6 | 1.1 | 7.2 | 1.5 | 5.6 | 1.6 |

We find a correlation between the diffusion coefficient and the complex mass, while we do not observe any correlation between the diffusion coefficient and the number of interactions of the analyzed proteins (Supplementary Fig. 3). However, since SbRD yields diffusion values corrected for confinement effects, we obtain a new and improved correlation between diffusion and complex mass, with the diffusion coefficient scaling as *D* = *αM*^−06^. Consequently, the proposed correlation between diffusion and perceived viscosity *(η)*^16^ changes *to η = αM_complex_^0.^*^27^ (Supplementary Fig. 3).

### SbRD versus SMdM analysis of the diffusion coefficient at the cell poles

Diffusion measured near the cell boundary and in the cell pole regions of rod-shaped bacteria appears slower than in the cell center due to confinement effects ^16, 38, 56–58^. We recently showed ^16^ that the ratio between the diffusion coefficient at the cell poles and cell center is lower for SMdM data than for simulated data, where particles are treated as mathematical points moving of random motion. This indicates that the slowdown observed in cells must be due to some physiological effects, such as increased crowding in the polar region, possibly due to aggregation of old or damaged proteins, the presence of the translation machinery, or dynamic structures generating over- and undercrowded regions ^16^. One of the key unanswered questions about the diffusion measured at the cell poles is: how much of the observed slowdown is due to confinement and how much is due to physiological effects? It is not possible to decouple these effects by SMdM or other single molecule microscopy techniques. The ratio between the diffusion coefficient at the cell poles and the cell center obtained by SbRD correlates linearly with the ratio obtained by SMdM (Supplementary Table 1). This is not observed in simulated cells (Fig. 7A). Therefore, the slowdown observed at the poles cannot be attributed solely to the effect of confinement. We compared the ratio between the diffusion at the cell poles and the diffusion at the cell center obtained via SbRD with the one obtained via SMdM for all the analyzed cells clustered as billiards. We observe a *D_pole_*/*D_center_* ratio of 0.74 ± 0.13 for SMdM and 0.80 ± 0.16 for SbRD (Fig. 7B). We analyzed the difference with a Mann-Whitney U rank test for non-normally distributed data and obtained a *p- value <<* 0.01. We therefore conclude that about 20% of the previously observed 25% slowdown at the cell poles can be attributed to confinement effects.

**Fig. 7.**
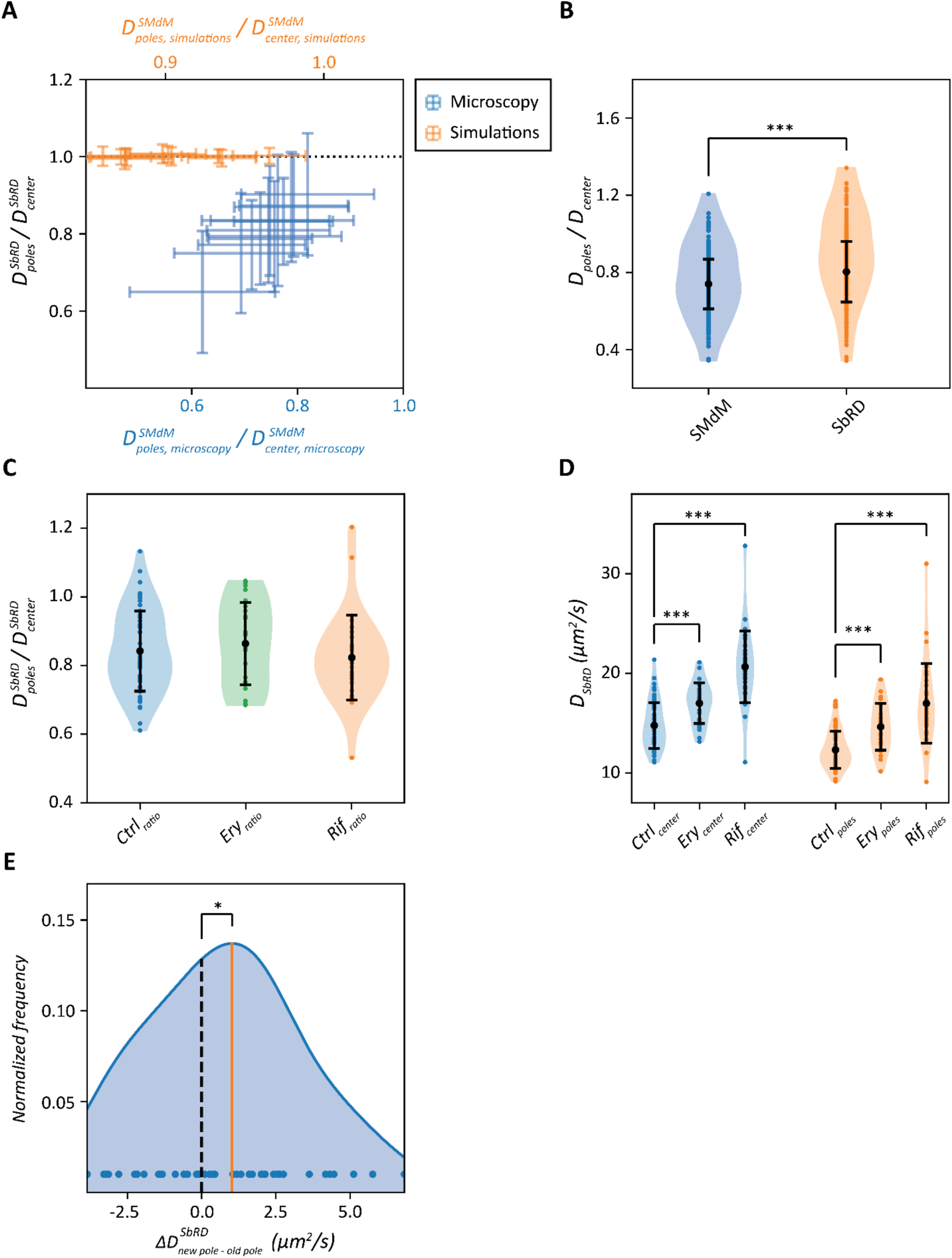
Cell pole analysis of diffusion. **(A)** Comparison of the ratios of the diffusion coefficients at the poles and center of the cell for data analyzed by SbRD and SMdM. The black dotted line shows the case when the diffusion at the poles and center is equal. For simulated data (orange) the ratio obtained with SbRD is equal to 1 for each protein. For microscopy data (blue) we find a positive correlation between the ratio obtained by SMdM and SbRD. (B) Ratios of diffusion at the poles and the cell center obtained by SMdM and SbRD. (C) Ratios of diffusion at the poles and cell center by SbRD for control cells, and cells treated with 250 µg/ml of erythromycin or 500 µg/ml rifampicin. **(D)** Diffusion coefficients at the cell center (blue) and poles (orange) for control, erythromycin- and rifampicin-treated cells. (E) Distribution of differences in diffusion coefficients between the newly formed cell pole and the old cell pole. The orange line represents the average of the distribution, the black dashed line is the zero. All curves are obtained via kernel density estimation. Statistical significance is indicated with asterisks.

### Effect of Rifampicin and Erythromycin on diffusion at the cell poles

The slowdown in diffusion at the cell poles observed via SbRD can be due to: (i) the presence of polysomes, large structures composed of several ribosomes bound to the same mRNA molecule; (ii) clusters of aggregated or damaged proteins that hinder the mobility of other molecules; (iii) or the presence of unknown substructures. To test the first hypothesis, we treated *E. coli* cells expressing mEos3.2 with the antibiotics rifampicin and erythromycin, which disrupt the polysomes via different mode of actions. Rifampicin inhibits DNA-dependent RNA biosynthesis by inhibiting the bacterial RNA- polymerase ^59, 60^, which leads to rapid RNA depletion, particularly of mRNA ^61, 62^, while erythromycin inhibits the assembly of the large ribosomal subunits 50S ^63^. Hence, the use of these antibiotics should make the diffusion coefficients at the poles and middle of the cell similar if the polysomes would form a major hindrance for the mobility of mEos3.2. However, we do not observe a significant change in ratio of the diffusion coefficients (Fig. 7C), with values of 0.84 ± 0.12, 0.86 ± 0.12 and 0.82 ± 0.12 for untreated cells, cells treated with erythromycin and cells treated with rifampicin, respectively. This is confirmed by the Mann-Whitney U rank test for non-normally distributed data. To further investigate the effect of the antibiotics, we analyzed the absolute values of diffusion. For cells treated with erythromycin, the diffusion at the cell center and at the cell poles show a moderate increase compared to the control sample, while for cells treated with rifampicin the increase is much larger (Fig. 7D). For the cell center regions we observe diffusion coefficients of 14.76 ± 2.30 µm^2^/s, 16.99 ± 2.04 µm^2^/s and 20.65 ± 3.61 µm^2^/s, and for the cell poles 12.31 ± 1.87 µm^2^/s, 14.62 ± 2.35 µm^2^/s and 16.98 ± 3.99 µm^2^/s, for cells untreated, treated with erythromycin and treated with rifampicin, respectively; the Mann-Whitney U rank confirms these findings (*p-values* << 0.01). We tentatively conclude that the overall faster diffusion in the presence of the antibiotics is the result of a lower viscosity due to the depletion of mRNA, which is most pronounce upon rifampicin treatment.

### Analysis of diffusion at the cell poles indicates asymmetry that correlates with aging

We then analyzed the microscopy data for differences between the cell poles (Supplementary Fig. 4). We reasoned that differences in aging of the two poles could lead to differences in diffusion, especially because misfolded and aggregated proteins tend to accumulate at the old pole ^19–21^. We acquired SMdM data of newly divided cells by selecting fields of view with cells that had just completed the division process (Fig. 5B, 5C), and we subsequently applied SbRD to the datasets. We determined the diffusion coefficient corrected for confinement effects of mEos3.2 in each cell individually and find that the diffusion at the old pole is significantly slower than at the new pole. The ratio between the diffusion at the new cell pole and the cell center is 0.86 ± 0.15, while *D_old-pole_/D_center_* is 0.80 ± 0.13 (Supplementary Fig. 5). With the assumption that the new cell pole enables on average faster diffusion than the old cell pole, we subtracted *D_old-pole_* from *D_new-pole_* for each cell. We obtained a distribution of residual diffusion coefficients with a mean higher than zero (Fig. 7E). We performed a one-sided Wilcoxon signed-rank test to confirm whether the observed difference was significant, and obtained a *p-value <* 0.05. These findings confirm the hypothesis that aging influences the structure of the cytoplasm at the poles of *E. coli,* causing macromolecules at the older cell pole to diffuse slower than at the new one.

## Discussion

We developed a new method to obtain diffusion coefficients that are not affected by confinement effects and bias by 2D modeling of a 3D motion. This method is key not only for measuring lateral diffusion in small compartments, but also for diffusion in proximity of boundaries, such as the plasma or organellar membranes. Despite the enormous advancements offered by SMdM in probing molecule motion compared to other methods, the values obtained near boundaries are affected by the effect of confinement. Our newly developed method SbRD allows examining more precisely regions of the cell that are small and or geometrically more complex.

### A method to reconstruct confinement-corrected diffusion coefficients in small cells

The main advantage of the SbRD method is that the shape of the analyzed compartment does not limit the analysis. In fact, compartments of any shape, that can be visualized and reconstructed via triangularization, can be used to recreate an identical virtual compartment. In this way, our method allows reconstructing unbiased diffusion in heterogeneous compartments such as those in eukaryotic cells.

We show by simulations that lateral diffusion measurements performed in small compartments, such as the prokaryotic cell, are bound to underestimate the diffusion coefficient, not only in the regions near the boundary but also in the cell center. The confinement effect in the cell center is mostly due to 2D observation of a motion in 3D. Using the here developed tool for cell clustering, we precisely detect spatial information such as radius, length, center and orientation angle of a cell. We use this information to reconstruct cells with an assumed spherocylindrical shape in Smoldyn ^4S^. Recursive diffusion simulations then yield the diffusion coefficient corrected for the confinement effect and the spatial component of motion. This approach led us to (more precisely) estimate the dependence of the diffusion coefficient on the complex mass of the diffusing species and infer from the data the viscosity perceived by a molecule of given molecular mass.

We observed a linear dependence of the ratio between the diffusion at the cell poles and the cell center for data acquired via SbRD and via SMdM, as shown in Fig. 7A, indicating that the observed slowdown in diffusion at the cell poles can be attributed to physiological effects, and not solely to confinement.

### Asymmetric diffusion at the cell poles correlates with aging

We previously obtained indications that the diffusion at the cell poles is slower than in the center of *E. coli* cells ^16^. We now determine precisely how much slower the diffusion is at the cell poles, and we show that disassembly of polysomes and depletion of mRNA by antibiotic treatment do not affect the differences in diffusion between the poles and the center of cell. In fact, the diffusion coefficient increases by a similar percentage in each region of the cell, suggesting that the antibiotic treatment has decreased the overall viscosity of the cell.

Moreover, we were able to precisely detect dividing cells and the new cell poles, and we show that the ones at the division site exhibit a faster diffusion compared to the old cell poles. The diffusion coefficients at the new and old pole are 86% and 80%, respectively, of the value measured at the cell center. In eukaryotic cells aging is accompanied by an increased cytosolic crowding ^64^. Here, we hypothesize that relative slowdown of diffusion at the old pole is consistent with an increase in crowding and an indication of aging in *E. coli*.

Previous studies suggested the possibility of accumulation of aggregated proteins at the cell poles of *E. coli* as a possible cause for the observed aging effects ^19^. In the recent study from Łapińska et al. ^45^, however, the authors do not observe protein aggregation in the form of inclusion bodies and they argue that proper techniques for the investigation of the structure of cell poles are lacking. Here, we developed a method to more precisely assess the structure of the cytoplasm in the cell pole regions. We also do not see indications of protein aggregation such as inclusion bodies in our dataset. We conclude that the slower diffusion observed at the old cell pole is an indication of the presence of partially aggregated misfolded macromolecules.

The *E. coli* cell cycle is divided in three different phases: the B period, which goes from the cell birth to the beginning of DNA replication; the C period, which goes from the beginning to the termination of DNA replication; and the D period, which represents the time between the termination of DNA replication and cell division ^65–66^. During the D period, the two chromosomes segregate to two different parts of the cell, and a period of protein synthesis is necessary for completing the cell division process. If accumulation of static or semi-static structure is a sole characteristic of the cell poles, then the newly formed cell pole should exhibit a diffusion value close to the one measured at the cell center. Since we observe a slower diffusion at the new cell pole (albeit less slow than at the old pole) compared to the one measured at the cell center, we conclude that the accumulation of damaged or aggregated protein is not a specific characteristic of the cell pole, rather it is a consequence of steric hindrance exerted by the nucleoid, which causes accumulation of structures in nucleoid-free regions of the cell. Based on the currently available literature ^19, 44, 45, 67^, we hypothesize that the observed slowdown is maintained throughout the cell cycle, and that it is possibly passed down from mother cell to daughter cells, where one of the daughter cells will inherit the “slow pole“ from the mother, while the other will inherit the “fast pole“ (Fig. 8).

**Fig. 8.**
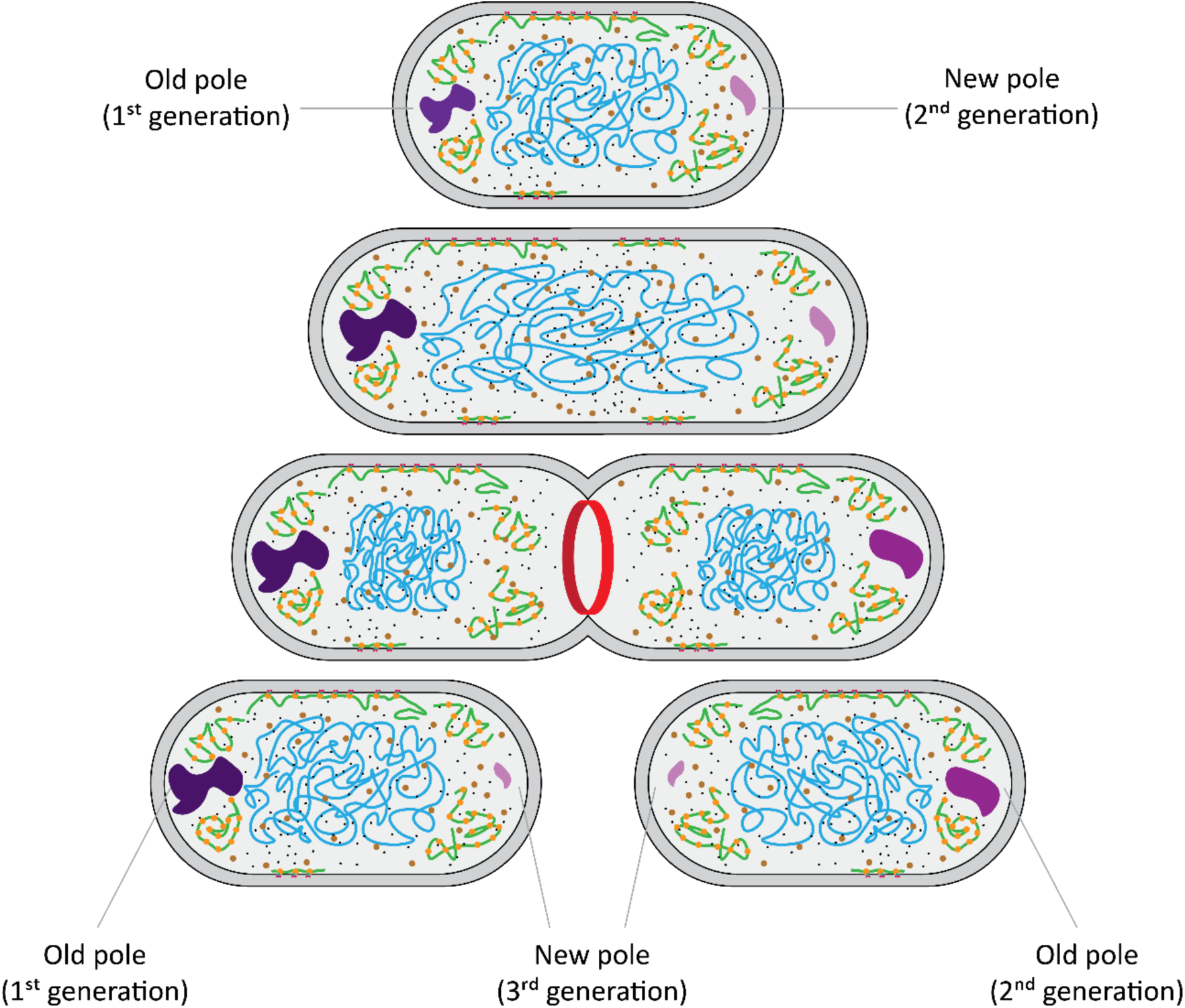
Accumulation of aggregated structures at the cell poles and correlation with aging. A cell with an old pole, with slower diffusion, and a new cell pole, with faster diffusion is shown at the top. As the cell cycle progresses this difference is maintained. When the cell divides and the septation ring forms (red), the two daughter cells will inherit the two cell poles. One of the cells will have the oldest pole as its old pole, while the other will have the new pole of the mother as its old pole.

### Concluding remarks

We developed Simulation-based Reconstructed Diffusion (SbRD) to determine diffusion coefficients in any compartment, corrected for confinement effects and by the motion of particles along the *z*-axis. We applied the technique to a previously recorded dataset. Using SbRD, we obtain a more precise correlation between the diffusion coefficients and the complex mass of the diffusing species, from which we infer the perceived viscosity for proteins diffusing in the cytoplasm. We recorded new single-molecule displacement datasets to characterize the slower diffusion at the cell poles, and to determine differences in confinement-corrected diffusion at the old and new pole. We correlate slower diffusion at the old poles with aging of the cells. We argue that our method and analyses provide new possibilities for investigating the mechanism of aging of bacteria and other types of cells.

## Materials and methods

### Live-cell single-molecule microscopy

Media preparation, cell culturing, measurements setup and live-cell imaging was performed as described ^16^. Briefly, for each experiment we started a pre-culture of *E. coli,* bearing a pBAD plasmid for the expression of mEos3.2, by scratching a glycerol stock with a sterile inoculation loop and dipping it in a 14-mL plastic culturing tube containing 3 mL of LB medium, prepared following the formula of 10/10/5% (w/v) in MilliQ of NaCl, tryptone (Formedium), and peptone (Formedium), respectively, and supplemented with ampicillin (100 µg/mL). We incubated the pre-culture overnight at 30°C, with shaking at 200 rpm. On the following day we transferred 30 µL of the LB pre-culture into 3 mL of MOPS-buffered minimal medium (MBM), prepared following the formula in ^68^, supplemented with 0.1% (v/v) glycerol and ampicillin (100 µg/mL). Cultures were incubated overnight at 30°C, with shaking at 200 rpm. The next day cells were diluted to a final OD_600_ of 0.05 to 0.08 into prewarmed MBM containing 0.1% (v/v) glycerol, ampicillin (100 µg/mL) and 0.1% (w/v) L-arabinose, and incubated the at 30°C, with shaking at 200 rpm for 4 to 6 hours before microscopy experiments. Right before the measurements, the cultures were spun down in a tabletop centrifuge and concentrated three times in the growth medium.

To ensure a constant temperature of the microscope during the imaging process, the instrumentation was turned on 4 to 5 hours before the measurement, to minimize the *xy* drift of the samples. Cells were imaged on a clean, non-functionalized high-precision glass slide {specs, manufacturer), previously sonicated in 5M KOH for 45 minutes and then rinsed 10 times with MilliQ, followed by a drying process via pressurized air. Immobilization of the samples was achieved by depositing 5 µL of concentrated cell suspension on the glass slied and then pressing the cells against the glass surface with solidified agarose pads having the same composition of the MBM medium with a final concentration of agarose of 0.75% (w/v), formed inside a polydimethylsiloxane (PDMS) chamber.

Once the cells settled, we selected an area of our field of view to perform the measurements. We adjusted the focus and the laser beam angle to obtain the highest number of foci, which resulted in the beam angle slightly below that of the critical angle for total internal reflection [highly inclined and laminated optical sheet microscopy ^69^]. The camera and the laser were then synchronized in the stroboscopy mode, with illumination pulses necessary to first photoconvert and then detect mEos3.2 every 1.5 ms ^16, 35^. For a detailed overview of the scripts used for managing the microscope, we refer to our code ^70^.

Erythromycin treatment was performed for 1 hour after cells reached an OD_600_ of 0.12-0.15 on the day of measurement, by adding erythromycin to the cell culture to a final concentration of 250 ng/µL. Agarose pads were supplemented with the same erythromycin concentration.

Rifampicin treatment was performed as described for erythromycin, using a final concentration of rifampicin of 500 ng/µL. Agarose pads were not supplemented with rifampicin, as this influenced the photoconversion of mEos3.2.

To analyze dividing cells, we visually inspected different fields of view in every sample, and selected areas in which a pair of obviously dividing cells were observed. Dividing cells were then analyzed separately during data analysis.

### Cell clustering and detection

The first step in our data analysis pipeline is represented by the detection and clustering of the cells in the imaged field of view. First, the whole analyzed field of view was converted into a 2D histogram in which every bin represents a certain number of fluorescent points. The background value was calculated by taking the median value of the bins. Only the fluorescent points belonging to bins having a value higher than the background value plus one standard deviation were kept and used to create point clouds. For each point cloud the eigenvectors were obtained through the calculation of the covariance matrix, which allowed calculating the angle between the first eigenvector and the x axis. From this, an appropriate rotational matrix was applied to the *xy* coordinates of the point cloud, to align its major axis parallel to the x-axis. This allowed obtaining a first set of features of the point cloud, namely the length, the width, the rotation angle and the center, which we used to describe a billiard encompassing the point cloud. Shape refinement was obtained by fitting an improved billiard around the point cloud via maximum likelihood estimation. The fitting was performed by applying the following assumptions: (i) the fluorescent molecules are uniformly distributed throughout the cells. Since the cells are spherocylinders imaged in two dimensions, the number of molecules observed is directly proportional to the thickness of the cells; therefore the boundary areas are less populated than the center; (ii) every observed point is equally likely as any other to be due to random noise; (iii) the probability of a point being random noise is equal to the probability of a point being a fluorescent molecule. From these three assumptions we obtain the following probability mass function for each particle (Eq. 9):

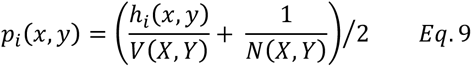

Where Λ/ is the number of particles *(X,Y)* inside the spherocylinder, *V* is the volume of the spherocylinder, which is modeled based on all the points *(X,Y)* within the spherocylinder, and *h_x,y_* is the thickness of the spherocylinder at the *xy* coordinate of the detected point. All these parameters depend on the size of the spherocylinder, which is described by its length, its radius, its center and its rotation angle. Therefore, these are used as fitting parameters to identify the best spherocylinder describing the detected point cloud.

The identified cells are then filtered based on their shape, discarding cells that are shorter than 0.65 µm (possibly cells that are partially out of the field of view), or wider than 1.5 µm (possibly noise or drift). Cells having a length bigger than 3 µm were automatically reanalyzed as dividing cells. In case of overlapping billiards for dividing cells, the intersection points were identified by calculating the intersection of the two semicircles describing the two adjacent cell poles. From this, it was possible to calculate the volume of the two spherical sections of the neighboring cell poles. Since any of the observed points could belong to either cell, the final volume used to calculate the probability density had to be adjusted by adding the intersection volume (Eq. 10):

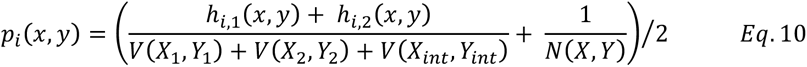

Where *N* is the number of particles *(X,Y)* inside the two spherocylinders, *V(X_1_*,*Y_1_)* is the volume of the first spherocylinder, *V(X_2_,Y_2_)* is the volume of the second spherocylinder, *V(X_int_,Y_int_)* is the volume of the intersection between the two spherocylinders, *h_1_* is the thickness of the first spherocylinder at the *xy* coordinate of the detected point, and *h_2_* is the thickness of the second spherocylinder at the *xy* coordinate of the detected point.

Finally, the identified cells are visually inspected and discarded if not suitable for analysis (e.g. dividing cells not correctly identified, for which the total length is shorter than 3 µm). The fitted spherocylinders are then used to create clusters from the point clouds. The data analysis is then performed on each cluster separately, ignoring the points that are not included in the cluster. More information on the cell detection and clustering can be found on our code ^70^.

### SMdM analysis

SMdM analysis was performed as described previously ^16^, with the exception that cell clustering was performed prior to peak pairing. Briefly, we recorded several consecutive movies for each field of view and paired the observed localizations from the two consecutive frames of the stroboscopic illumination pattern. For single-molecule analysis, we used the STORM-analysis package developed by the Zhuang laboratory, which is included in the 3D-DA0ST0RM program for peak detection ^71^. After a full movie was analyzed, the localizations were corrected for *xy* drift.

All the detected peaks in each field of view were used for clustering and for finding the shape of the spherocylinder that best describes the shape of the cell, as reported in the section above.

Displacements were obtained from all the peaks belonging to a single cell, by pairing localizations from the two consecutive frames of the stroboscopic illumination pattern. We set a maximum distance of 600 nm between any two peaks to be paired: the distance is then used to find all possible peak pairs for each couple of frames. To obtain a displacement, we match each peak in the first frame of the couple with all the peaks falling within a radius of 600 nm in the second frame. This procedure is repeated for all frame couples of each field of view. A hard filter based on the number of detected displacements was then applied, discarding cells with less than 2000 or more than 20000 displacements, as described ^16^.

A pixel map with pixel size of 100 nm^2^ was obtained for each cell, with every pixel containing the information of all the peak pairs for which the starting position is located inside the pixel itself. Each pixel of the map containing a minimum number of displacements (set to 10 in our study) was then fitted using a modified two-dimensional probability density function (PDF), which accounts for a linear background effect *k,* which can be caused by an ambiguity in the assignment of peak pairs ^16^ (Eq. 11):

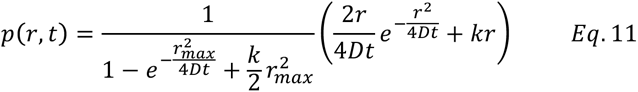

Since *t* is known, as it represents the time between two stroboscopic laser pulses, the PDF was fitted on the detected displacements *r,* using the diffusion coefficient *D* and the background value *k* as fitting parameters. Displacements were detected using the MLE clustering method (see section above) for SbRD. Displacements were detected both using the MLE clustering method and Voronoi clustering when comparing the two clustering methods using SMdM.

Given our advanced cell detection method, we could perform an accurate identification of the different cell regions, namely the cell poles and the cell center. Fitting a billiard to detect the shape of the cells allows obtaining precise information about their length and their radius, which in turn allow identifying the cell pole regions and the central region of the cell. All the displacements belonging to the same region were then used to perform the fitting using equation 10, yielding information about the diffusion coefficient in the different regions of the cell ^16^.

The dependence of the diffusion coefficient on the complex mass was fitted using a power law relationship *D = αM_complex_^β^,* where *M_complex_* is the complex mass and *α* and *β* are fitting parameters ^16^. Fitting was performed using the function *curve_fit* included in the SciPy library^72^.

### Smoldyn simulations

Simulations were performed using the software Smoldyn ^46^, as described ^16^. A diffusion coefficient and a time-step length are used as input for the simulations, together with the total simulation time. At every time step, Smoldyn randomly selects a step length from a normal distribution having as mean the squared mean squared displacement calculated from the input diffusion coefficient, as well as a random direction in the *xyz* space for each particle. These values are used to simulate the motion of every particle in the system at every time step, until the total simulation time is reached. In our simulations we used a time step of 0.1 ms and a total simulation time of 2 seconds. Particles every 15 steps (1.5 ms) were then paired together in displacements, the results were benchmarked against the microscopy data.

We used Smoldyn to generate two separate datasets. First, we simulated the motion of particles using input diffusion coefficients ranging from 0.01 to 110 µm^2^/s in a spherocylinder having length and width of 2.25 and 0.9 µm, respectively, as these reflect the average cell size observed in our previous work ^16^. We then generated a second dataset using input diffusion coefficients ranging from 1 to 20 µm^2^/s in a spherocylinder with a length ranging from 1.4 to 2.9 µm and width ranging from 0.6 to 1.5 µm, always keeping the ratio between length and width higher than 2 and lower than 4 to reproduce actual dimensions of *E. coli*.

We analyzed the output of our simulation using the same approach adopted for SMdM measurements and compared the results with those obtained by SbRD (see section *SbRD analysis).* The equation used for simulated data does not account for linear background correction (Eq. 12):

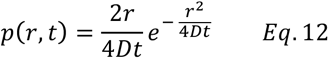

More information about the simulations can be found on our code ^70^.

### SbRD analysis

Simulation-based Reconstructed Diffusion (SbRD) was applied by using Smoldyn simulations on SMdM analyses. A collection of points representing a cell, either coming from a Smoldyn simulation or from a microscopy experiment, is analyzed using SMdM (see section *SMdM analysis).* This analysis provides a pixel map of the cell, in which for each pixel the diffusion coefficient, as well as the *xy* start and end positions of each displacement, are known. For microscopy measurements, the new cell clustering method allowed us to determine the length and radius of the spherocylinder in which the diffusing particles are confined, which could be modeled in Smoldyn. In the case of dividing cells, the septum is modeled as a reflecting wall passing through the intersection points of the two billiards, encompassing two separate cells, and parallel to the *z* axis. In the case of data from Smoldyn simulations, the length and radius of the spherocylinder are known. These information were then used to start a recursive simulation in Smoldyn by placing a number of particles equal to the number of displacements in their respective *xy* starting position inside the pixel, with the *z* position randomly assigned to each particle inside the spherocylinder, which in the case of microscopy data was modeled as described in *Cell clustering and detection.* A simulation lasting for 1.5 ms, with simulation steps of 0.1 ms is started, using as input diffusion coefficient the value of the pixel obtained via SMdM (see section *Smoldyn simulations).* The output of this simulation is then used to perform a fitting using equation 11, with *D* as fitting parameter. The squared difference between the output diffusion obtained via simulations and the diffusion obtained via SMdM is then calculated. The program then recursively iterates the simulation process until such squared difference reaches a minimum. The input diffusion coefficient used to obtain the output diffusion coefficient that minimizes the squared difference is then regarded as the real diffusion coefficient of the pixel. This process is then repeated 10 times for each pixel to account for the randomness introduced by Smoldyn ^46^ in the choice of the step length and the direction of motion, as well as for the randomness introduced in the placing the particle along the *z*-axis. The process is then repeated for every pixel of the original SMdM map, from which an SbRD map is obtained. The pixel-by-pixel differences between the SbRD map and the SMdM map are used to construct a difference map. More information about the SbRD analysis can be found on our code ^70^.

### Statistical analysis

All statistical analyses were performed using the Python package *stats* from the SciPy library ^72^. Shapiro-Wilk test for normality ^73^ was used to check whether the data are normally distributed, using a level of confidence of 5%. The test assumes the null hypothesis for data that are normally distributed. Therefore, if the obtained *p-value* is lower than 0.05 the null hypothesis is rejected and the data are assumed to be non-normally distributed. Non-normally distributed datasets are visualized via kernel density estimation.

The Mann-Whitney U rank test^74^ was used to test whether the means of two non-normally distributed datasets are equal. In the case of comparing means of datasets, *i.e.* when no prior assumptions are made and no precise outcome is expected, such as in the case of comparing the diffusion coefficients of cells treated with antibiotics, a two sided test was performed. In the case of comparing means of datasets in which a specific outcome was expected, such as in the case of comparing the diffusion coefficient of the two different cell poles, a one sided test was performed.

The Wilcoxon signed-rank test^75^ was used to test whether the median of a dataset coming from paired measurements is significantly different from zero. This test was used to assess whether the difference in diffusion between the new cell pole and the old cell pole was significantly higher than zero, therefore it was conducted as a one sided test. Statistical significance in pictures are indicated with 1 asterisk (*) for *p-value < 0.05,* 2 asterisks (**) for *p-value < 0.01* and 3 asterisks (***) for *p-value < 0.001*.

## Author contributions

Conceptualization: L.M. and B.P.

Experimental design: L.M. and B.P.

Mathematical modeling: L.M. and HJ.K.

Microbiology: L.M., D.S.L. and W.M. Ś.

Data acquisition: L.M., D.S.L. and W.M. Ś.

Python scripting: L.M.

Simulations: L.M.

Data analysis: L.M.

Statistical analysis: L.M.

IT supervision: M.P.

Project supervision: B.P.

Writing – original draft: L.M.

Writing – review and editing: B.P.

## Data and materials availability

All data needed to evaluate the conclusions in the paper are present in the paper and/or the Supplementary Materials.

